# Exploring the effect of Relative Language Distance on Bilingual Brain Structure – a cross-sectional VBM study

**DOI:** 10.1101/779751

**Authors:** Keerthi Ramanujan

**Affiliations:** Division of Speech and Hearing Sciences, Faculty of Education, The University of Hong Kong, Hong Kong; Centre for Cognition and Decision Making, Institute for Cognitive Neuroscience, HSE University, Russian Federation

**Keywords:** bilinguals, language distance, typological similarity, language control, brain structure, VBM

## Abstract

It is known that bilinguals’ perpetual need for language control influences their brain structure in significant ways. But bilinguals’ language control needs are themselves influenced by key dimensions of the bilingual experience – variation in the age of bilingualism-onset, relative language proficiency, exposure and immersion has indeed been shown to have differential effects on bilingual neurostructural profiles. An under-studied dimension of bilingualism that could also generate differing bilingual language control needs is the extent of similarity between bilinguals’ language pairs, referred to in the present study as Relative Language Distance (RLD). The goal of the present study was to explore whether the experience of managing “close” and “distant” languages has any impact at all on bilingual brain structure. To this end, exploratory morphometric analysis of grey matter volumes was carried out on three groups, all very similar in their bilingual profiles except for the factor of RLD – high-distance Cantonese-English (hd-CE), intermediate distance Hindi-English (id-HE) and low-distance Dutch-English (ld-DE) speakers. The results after controlling for contribution of other bilingual dimensions revealed significant bilateral putaminal volume differences between the groups that varied along the relative language distance gradient in the pattern of CE>HE>DE. This might be attributable to the differing articulatory control needs that variation in L1-L2 RLD generates. The present study reveals how the dimension of Relative Language Distance could impact bilingual brain structure.

## Introduction

Bilinguals are perpetually faced with the prospect of having to speak effectively in a chosen language despite the ever-present impediment of cross-linguistic interference (Kroll et al., 2015). They overcome this issue by relying on Language Control, a set of cognitive processes that facilitate adequate communicative competency in the desired language. Language control is implemented by regions in the prefrontal, parietal, cingulate cortices and the subcortical basal ganglia that together comprise the Bilingual Language Control (BLC) network (Abutalebi & Green, 2016; Calabria et al., 2018).

As a way of coping with the perpetual demand for language control imposed by the knowledge and use of two languages, adaptive neuroplastic changes can manifest in the BLC network (Green & Abutalebi, 2013). Several neuroimaging studies have reported adaptive structural capacity changes in bilingual brains (relative to monolingual ones) that are specifically attributable to dual-language use. Notably, the sites of such bilingualism-associated structural neuroplasticity are often found to occur within the BLC network. Grey matter volumetric changes have been reported in the inferior parietal lobules (IPL) (e.g., left IPL: Mechelli et al., 2004; bilateral IPLs: Abutalebi et al., 2015b), left inferior frontal cortex (IFC) (e.g., Klein et al., 2014), the anterior cingulate cortex (ACC) (e.g., Abutalebi et al., 2012; 2015a) and basal ganglia (e.g., left Putamen: Abutalebi et al., 2013b; left Caudate: Zou et al., 2012). These BLC regions have also been reported as the loci of grey matter-related increases in studies of adult second language learning (e.g., left IFC & IPL: Legault et al., 2018; left IFC: Stein et al., 2010), and in studies involving specialized linguistic training, such as simultaneous interpretation (SI) (e.g., right IPL: Hervais-Adelman et al., 2017; left IFC: Martensson et al., 2012). These findings are not surprising. Both L2-learning and SI training are instances of language control needs getting amplified. In the process of becoming bilingual, monolingual learners come to need *inter*-language control which was previously not required. SI trainees though already bilingual, are learning to master a more extreme and precise form of language control in order to perform rapid and accurate translation. Taken together, such structural neuroimaging findings^1^ support the notion that observed structural changes in bilingual brains are in fact neuroplastic adaptations arising due to the persistent requirement for dual-language control (Bialystok et al., 2012; Perani & Abutalebi, 2015).

If neuroplastic structural consequences of bilingualism are believed to arise from the continuous reliance on language control resources, then factors modulating bilingual language control should also modulate bilingualism-associated structural changes. Indeed, key experiential dimensions of bilingualism are known to have an impact on bilingual neuroanatomical profiles, affecting particularly regions in the BLC network. This includes relative language proficiency (e.g., Abutalebi et al., 2013b; Mechelli et al., 2004), exposure and immersion (e.g., Abutalebi et al., 2015b; Pliatsikas et al., 2017), and Age of L2 acquisition/bilingualism onset (Klein et al., 2014; Mechelli et al., 2004). A dimension of bilingualism that has received considerably less attention is the relation between bilinguals’ language pairs. Considering that two entire linguistic systems, each with their own sets of morphosyntactic, phonological and orthographic features, coexist in the bilingual mind and brain, would the relative degree of similarity between them affect language control and in turn, affect the structure of the BLC network implementing it? This is the issue explored in the present study.

The extent of similarity between bilinguals’ spoken languages, referred to here as relative language distance (RLD), can have consequences for bilingual language control. Arguably, ‘closer’ the distance between a bilingual’s L1 and L2, greater the cross-linguistic conflict generated. As a result, bilinguals speaking relatively closer language pairs may require greater language control resources than their counterparts speaking linguistically distant language-pairs. For bilinguals whose languages are distant, perceptible language-distinguishing lexical features such as script (e.g. Latin letters vs. Arabic/Japanese/Hindi letters) and/or phonology (e.g. English phonemes vs Cantonese tonal phonemes), may even promote language-specific access to some extent (Gollan et al.,1997; Hoversten et al, 2015; Kroll et al., 2006), minimizing the extent of cross-linguistic conflict and thereby possibly reducing the load on language control resources. Since language control demands can vary with RLD of language pairs, how would sustained experience of managing such ‘close’ versus ‘distant’ language pairs affect bilingual brain structure? This question has not been well explored. A structural neuroimaging study by Abutalebi et al. (2015b) is the only one thus far to have shed some light on this matter. Results of this study revealed that increasing L2 proficiency in bilinguals with lifelong experience in speaking two linguistically close languages (Cantonese-Mandarin) was associated with greater GM volume changes in the left inferior parietal lobule (a BLC region) than for lifelong bilingual speakers of a distant language pair (Cantonese-English). This finding suggests a potential role for RLD in influencing bilingualism-associated adaptive neuroplasticity. The current study sought to explore this issue further, this time in healthy young adults.

The aim of this study was to explore whether the factor of RLD has any discernible impact on bilingual brain structure. Exploratory structural neuroimaging analyses were carried out to this end and involved three groups of early, high-proficient bilinguals with comparable bilingual experiences, age, education levels and socioeconomic statuses. Critically, the groups differed only with respect to their language profiles and therefore in the RLD of their spoken L1-L2 combinations. Based on the relative similarity of perceptible lexical-level features (script and phonology) of their L1-L2 pairs, Dutch-English (DE) participants were categorized as ‘low-distance (ld)’ bilinguals, Cantonese-English (CE) participants as ‘high-distance (hd)’ bilinguals and the Hindi-English (HE) participants as ‘intermediate-distance (id)’ bilinguals. It was predicted that structural differences, specifically in terms of grey matter volume, should exist within the BLC networks of the three bilingual groups who by virtue of variation in their respective RLDs have been experiencing varying language control needs. Considering the scarcity of previous empirical studies on the effects of RLD on BLC structure and function, specific predictions as to where exactly within the BLC network RLD-mediated effects may arise are not made. In any case, an affirmative result would lend support to the notion of RLD being an influential dimension of bilingualism capable of affecting bilingual brain structure.

## Materials & Methods

### Participants & Language Assessment

The study sample included 19 CE, 19 HE and 20 DE adult biliterate bilinguals who had also taken part in a functional imaging study (Ramanujan, 2019, *preprint*). All were right-handed with no known history of neurological and psychiatric disorders, with comparable age, years of formal education and socioeconomic levels (*p* >.01) (see Table 1). Moreover, without exception, all subjects were proficient non-immigrant speakers of their respective L1s (Cantonese/Hindi/Dutch) and were equally proficient in their L2 (English) which they acquired fairly early. Having spent a majority of their lives in regions where the L1 was indigenous and the L2 a prevalent foreign language of the region, all subjects at the time of the study had comparable exposure and immersion in dual-language contexts. For a more detailed description of the bilingual groups and language testing, please see Ramanujan, 2019, *preprint*.

**Table 1:**
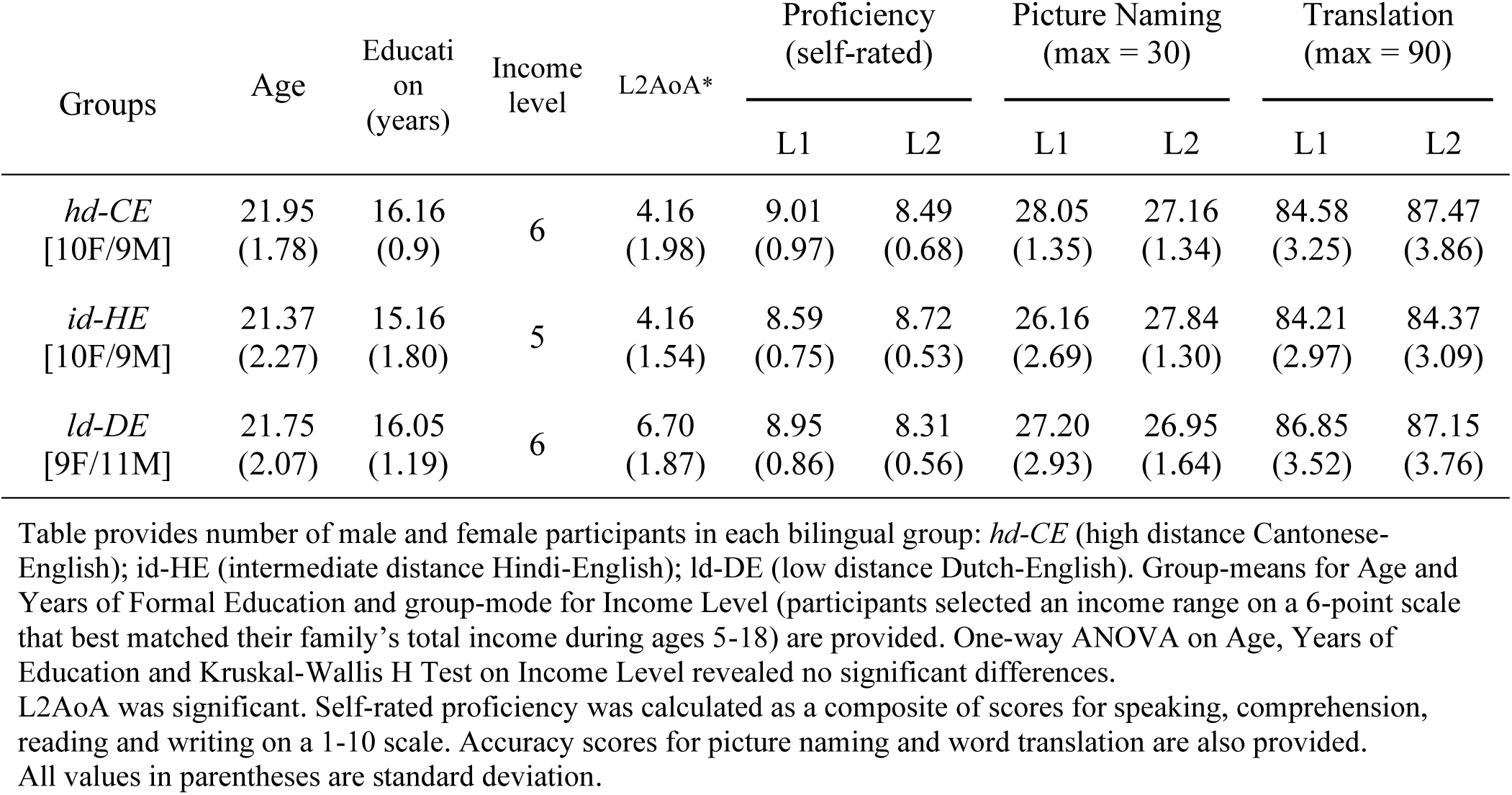
Demographic and Linguistic variables.

| Groups | Age | Education<br>(years) | Income<br>level | L2AoA* | Proficiency<br>(self-rated) |  | Picture Naming<br>(max = 30) |  | Translation<br>(max = 90) |  |
| --- | --- | --- | --- | --- | --- | --- | --- | --- | --- | --- |
|  |  |  |  |  | L1 | L2 | L1 | L2 | L1 | L2 |
| *hd-CE*<br>[10F/9M] | 21.95<br>(1.78) | 16.16<br>(0.9) | 6 | 4.16<br>(1.98) | 9.01<br>(0.97) | 8.49<br>(0.68) | 28.05<br>(1.35) | 27.16<br>(1.34) | 84.58<br>(3.25) | 87.47<br>(3.86) |
| *id-HE*<br>[10F/9M] | 21.37<br>(2.27) | 15.16<br>(1.80) | 5 | 4.16<br>(1.54) | 8.59<br>(0.75) | 8.72<br>(0.53) | 26.16<br>(2.69) | 27.84<br>(1.30) | 84.21<br>(2.97) | 84.37<br>(3.09) |
| *ld-DE*<br>[9F/11M] | 21.75<br>(2.07) | 16.05<br>(1.19) | 6 | 6.70<br>(1.87) | 8.95<br>(0.86) | 8.31<br>(0.56) | 27.20<br>(2.93) | 26.95<br>(1.64) | 86.85<br>(3.52) | 87.15<br>(3.76) |

Table provides number of male and female participants in each bilingual group: *hd-CE* (high distance Cantonese-English); *id-HE* (intermediate distance Hindi-English); *ld-DE* (low distance Dutch-English). Group-means for Age and Years of Formal Education and group-mode for Income Level (participants selected an income range on a 6-point scale that best matched their family's total income during ages 5-18) are provided. One-way ANOVA on Age, Years of Education and Kruskal-Wallis H Test on Income Level revealed no significant differences.
L2AoA was significant. Self-rated proficiency was calculated as a composite of scores for speaking, comprehension, reading and writing on a 1-10 scale. Accuracy scores for picture naming and word translation are also provided. All values in parentheses are standard deviation.

L2 AoA (age of Acquisition) varied significantly across groups (*F*_*2,55*_ *=* 12.99, *p* < .01, *η*^*2*^_*p*_ = 0.32). However, language assessment in L1 and L2 via a 90-item written word translation task and 30-item picture naming task (Abutalebi et al., 2007; Singh et al., 2017) revealed no significant proficiency differences in L2 across groups or between L1 and L1 within each group (see Table 1 for details) (*p* >.01). Additionally, subject’s self-reported L1 and L2 proficiency scores were also not significantly different (*p* >.01).

### Structural MR data acquisition

Structural data were acquired in a 3T Achieva Philips MR scanner (Philips Medical Systems, Best, NL) at the MRI centre of the University of Hong Kong.

An MPRAGE sequence (Magnetization Prepared Rapid Gradient Echo, TR=8.03 ms, TE=4.1 ms, flip angle = 8°, FOV = 250×250, matrix = 256, Sense factor = 1) was used to obtain high-resolution, 150-slice T1-weighted images of 1mm3 isotropic voxel size for each participant.

### Structural data pre-processing

MPRAGE T1 volumes of each subject were first visually inspected to check for gross field distortion and movement artefacts. No data was discarded at this stage. Then for each subject, the origin was manually set to correspond to the AC-PC (anterior commissure-posterior commissure) line. Following this, the structural volumes were processed using the Computational Anatomy (CAT12) Toolbox (ver. r1113, http://dbm.neuro.uni-jena.de/cat/, Gaser & Dahnke, 2016). The following preprocessing steps were implemented out for every subject with the default settings:

i. Pre-segmentation Denoising. Since noise is not equally distributed across MR images, a Spatial-Adaptive Non-Local Means (SANLM) denoising filter was implemented. This filter uses local information to adjust denoising filter strength to manage noise (Manjon et al., 2010).
ii. Local intensity transformation using Local Adaptive Segmentation (LAS) (Dahnke et al. 2012b). This procedure applies a local intensity transformation to correct any skewed local GM and WM estimations (Gaser & Kurth, 2016).
iii. Final Segmentation of volumes into Grey matter (GM), White Matter (WM) and Cerebrospinal Fluid (CSF) components. This routine was carried out using an Adaptive Maximum A Posterior (aMAP) technique in which tissue parameters are iteratively estimated and modelled locally at each site to generate data-driven custom tissue priors (Rajapakse et al. 1997). This reduces the dependency on a priori tissue probabilities while also accounting for local variations and inhomogeneity of GM intensity specific to each data.
iv. Regularization of segmented images using Affine regularization. In this step, individual CE data were registered to the ICBM (International Consortium for Brain Mapping) template for East Asian brains. HE and DE were registered to the ICBM European Brain template.
v. Normalization using DARTEL (Diffeomorphic Anatomical Registration through Exponentiated Lie algebra). In this final step, GM and WM segmentations were normalized to MNI space using a pre-existing DARTEL template (IXI555) derived from 555 healthy, normal adults (IXI database, http://www.braindevelopment.org). As the sample in the present study were not a special population and were similar in profile to the template population, this reduced the need to create a study-specific DARTEL template (see Gaser & Kurth, 2016).

Following the above preprocessing stages, a quality check for detecting any outliers within the groups was performed with CAT’s in-built ‘Check Sample Homogeneity’ option. Modulated and normalized GM images were used to compute group mean correlation and overall weighted image quality. No group’s mean correlation coefficient was lesser than 0.8. As no outliers were detected for overall weighted image quality, all subjects were retained for further analysis.

Segmented GM images of all subjects were subsequently smoothed using an 8mm FWHM (full width at half maximum) Gaussian kernel. Differences between the three bilingual groups were then investigated using conventional whole brain Voxel-based Morphometry (VBM) methods as well as Region of Interest-based Morphometry (ROI-BM).

### Whole brain VBM analysis

Following preprocessing, between-group whole brain differences were investigated. The smoothed, modulated and normalized GM images of each group were modelled as a Full Factorial design (1 factor *RLD* with 3 levels, CE, HE, DE). Total Intracranial Volume (TIV) in millilitres was computed for each subject by summing up native space global volumes of GM, WM and CSF to correct for any differences in brain volumes/size.

#### Step1: Estimating group differences at whole-brain level

To test for GMV differences (increases or decreases in GMV) across the groups arising due to RLD, a model with Education and Income level (proxy for Socioeconomic status), Age and TIV as nuisance covariates and L2 AoA as a potential confound was evaluated using an F-contrast. The results were assessed at p < .001 (uncorrected) at the voxel level with FWE (family-wise error) correction of p < .05 at the cluster level, with a spatial extent threshold of *k* ≥ 1000. Where voxels were significant but not clusters, SVC (small volume correction) was performed, set to FWE correction of *p* < .05. The contrast estimates at significant clusters were plotted to explore the direction of any detected difference for each group. Additionally, an F-contrast testing for any (positive and negative) correlations between GMV in any region in the whole brain and L2AoA was also evaluated at the same threshold.

#### Step2: Post-hoc Directional T-contrasts

To compare groups directly with each other the following T-contrasts (with the same covariates) were generated and evaluated at *p* < .001 (uncorrected) at voxel level with cluster-level FWE correction (*p* < .05) for a threshold of *k* ≥ 1000 voxels.

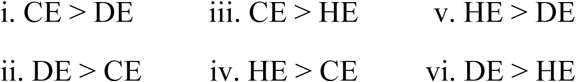

### Region-Based Morphometric (RBM) analysis

The logic of performing a whole-brain analysis and then following it up with a region-of-interest analysis is a standard practice in many VBM studies. However, the ROI analysis implemented in this study is slightly different from the conventional method. Instead of creating the usual 6-10mm spherical ROIs, atlas-based ROIs were used. The LPBA40 (LONI Probabilistic Atlas, Shattuck et al. 2008) was used to define ROIs. This atlas comprises probabilistic maps of 56 cortical and subcortical structures derived from 40 healthy volunteers (20 females) of ethnically diverse backgrounds similar in age and educational level to the subjects of the present study (see Shattuck et al., 2008). Following segmentation into GM, WM and CSF, a partitioning procedure was carried out in which, the probabilistic GM maps from the atlas are co-registered to subject’s native (non-normalized) space to retain any crucial subject-specific information. In this manner 56 custom-fitted ROIs were generated independently for each subject that closely matched the natural boundaries and volumes of cortical and sub-cortical regions.

#### Step1: Estimating group differences at ROI level

2^nd^ level GLM analysis was performed using the different atlas-based ROIs in CAT12. The same model used in the VBM analysis was evaluated using a F-contrast. This would test for any differences within all defined ROIs across population groups after removing effects of nuisance covariates and potential confounds. Thus, this type of RBM analysis, unlike the conventional ROI analysis, does not involve data-averaging which could possibly eliminate differences in voxels within a small region (Garcia-Penton et al., 2016) Also, while providing increased inferential power afforded by the likes of conventional ROI analysis and Small Volume Corrections (SVC), this method does not require *a priori* selection of regions of interest and in this way is similar to a data-driven, exploratory VBM approach.

Given the much smaller search volume, the RBM results were evaluated at p < .05 (uncorrected) with FDR (False Discovery Rate) correction for multiple comparisons. An F-contrast testing for any region-specific GMV correlations with L2AoA was also assessed at the same level of significance.

#### Step2: Post-hoc Directional T-contrasts

To compare groups directly with each other for the same sets of atlas-based volumetric ROIs, the same six T-contrasts as used for the whole-brain VBM analysis (with potential confounds modelled in) were evaluated at *p* < .05 at both uncorrected and FDR corrected thresholds.

### Brain-Behaviour Bivariate Correlations

The pre-normalization GM volumes were also extracted from the atlas-based ROIs of regions that both VBM and RBM identified and correlated with L2AoA and L1 and L2 Self-rated Proficiency scores.

## Results

F-contrast (main effects) results of VBM and RBM are presented first, followed by results of post-hoc directional t-contrasts between groups.

### Whole brain VBM Results

The Right Putamen varied significantly between the three bilingual groups, along with a cluster in the left temporal pole (see Figure 1 and Table 2). A supplementary analysis with the same covariates except for L2AoA also detected a significant difference in the right putamen (see Supplementary table S1). Test for any correlations between voxels in the whole brain with L2AoA did not yield any suprathreshold voxels or clusters. Contrast estimates plotted at the right putamen revealed that putaminal volumes were higher for CE and lower for DE (see Figure 1b).

**Table 2.** GM Differences between groups revealed through VBM.

| *Region* | *MNI co-ordinates (mm)* |  |  | *z-score* | *Cluster p*<br>*(FWE-corr)* | *k* |
| --- | --- | --- | --- | --- | --- | --- |
|  | *x* | *y* | *z* |  |  |  |
| **R Putamen** | **22** | **-1** | **10** | **4.03** | **.001** | **1122** |
| **L Superior Temporal Pole** | **-40** | **21** | **-32** | **3.70** | **.007** | **1048** |
| L Superior Temporal Pole | -39 | 24 | -38 | 3.36 |  |  |

**Figure 1.**
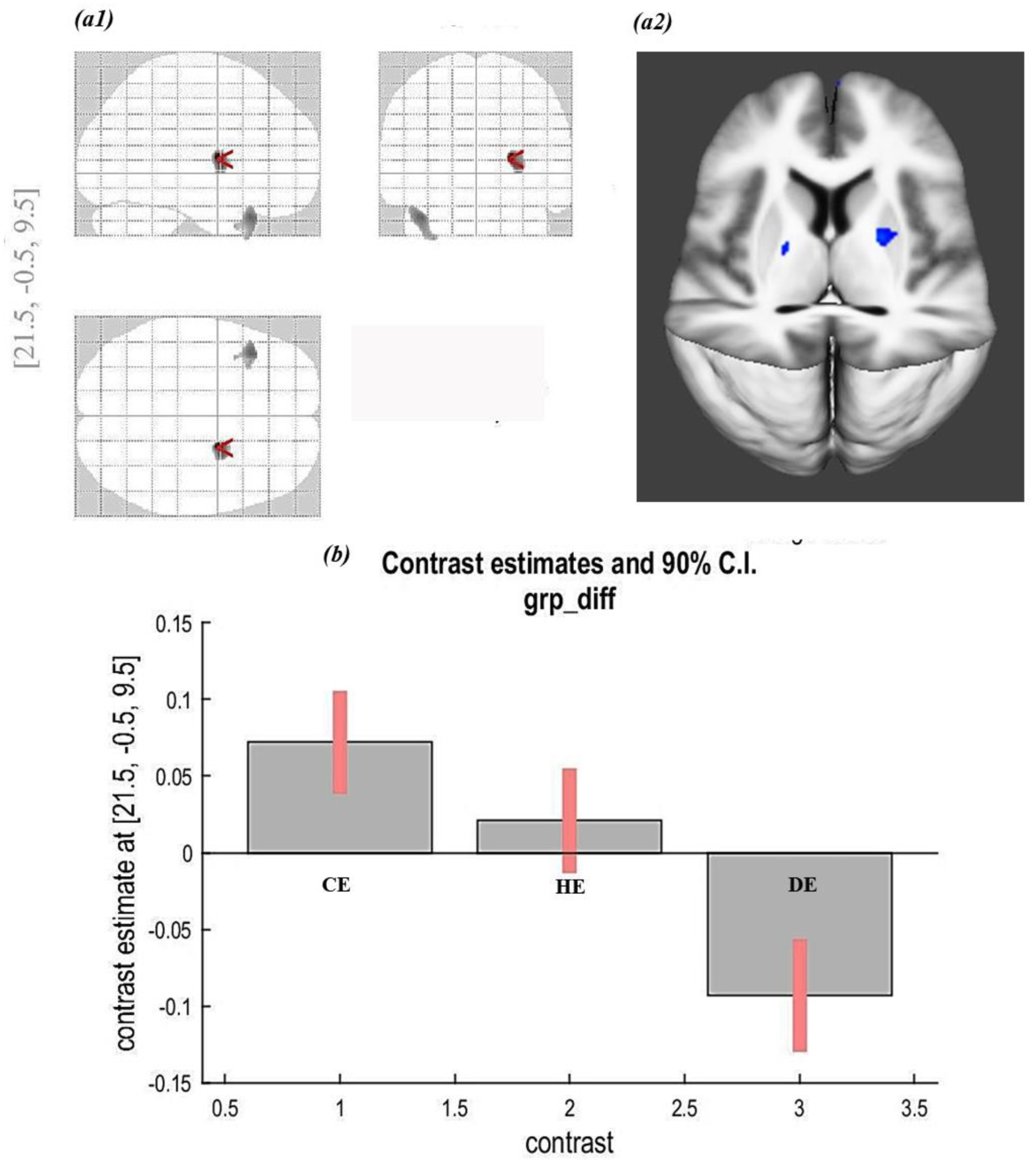
Whole-brain VBM Results: *(a1)* Between-group differences displayed on glass brain after removing confounding effects of education, socioeconomic status, age, TIV and L2AoA, revealing GM differences at the right putamen [22, -1, 10] ; *(a2)* F-contrast map showing suprathreshold voxels (in blue) in the right putamen [22, -1, 10] overlaid on a 3D rendering of the mean T1 image of the three bilingual groups. *(b)* Contrast estimates plotted at the same right putamen cluster showing direction of differences for the three bilingual groups.

### RBM Results

RBM results testing for any difference across groups, after controlling for education, SES, age, TIV and L2AoA, revealed differences in the bilateral putamen across the groups (see figure 2). Both right and left putamen were found to be significantly different even with FDR correction (*p* < .05) (see Table 3). A similar analysis with the identical covariates but without L2AoA, also revealed significant bilateral putamen differences (see Supplementary table S2). Test for correlations between any atlas-based ROIs and L2AoA did not yield any significant results.

**Table 3:** GM Differences between groups revealed via RBM.

|  | *p Value* |  |
| --- | --- | --- |
|  | uncorrected | FDR corrected |
| R Putamen | **< .001** | **0.03** |
| L Post Central Gyrus | 0.003 | 0.049 |
| L Putamen | **< .001** | 0.049 |
| L Superior Occipital Gyrus | 0.02 | - |
| L Middle Occipital Gyrus | 0.03 | - |
| L Lateral Orbital Gyrus | 0.03 | - |

Table provides uncorrected and FDR corrected p-values for ROIs that significantly varied across groups after controlling for education, socioeconomic status, age, TIV and L2AoA. ‘-’ indicates the ROI did not survive FDR correction.

**Figure 2.**
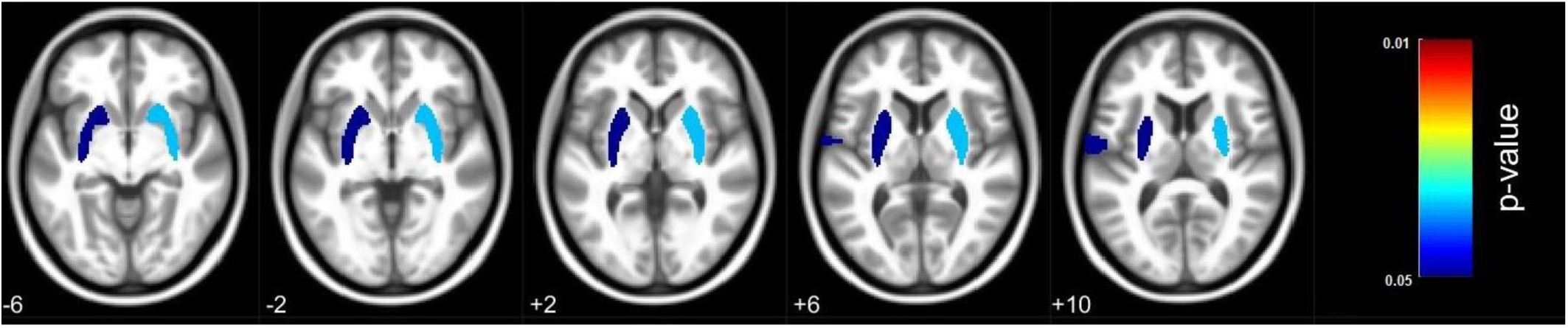
Whole-brain RBM Results: Differences between CE, HE and DE groups revealed through RBM analysis, displayed at FDR corrected levels. Left Putamen (dark blue) and Right Putamen (light blue) ROI’s GM volumes varied significantly across the three groups after correcting for Education, SES, Age, TIV and L2AoA.

### Post-hoc T-contrasts: VBM Results

Of the 6 directional post-hoc T-contrasts checking for whole brain VBM differences between the groups, the CE>DE contrast revealed significant suprathreshold voxels in the Right Putamen, Left Putamen and Left Superior Temporal Pole (see Table 4).

**Table 4:** VBM T-contrast results.

| Region | MNI co-ordinates<br>(mm) | | | z-score | Cluster $p$<br>(FWE-corr) | $k$ |
| --- | --- | --- | --- | --- | --- | --- |
| | $x$ | $y$ | $z$ | | | |
| *i. CE>DE* |  |  |  |  |  |  |
| **R Putamen** | **22** | **-1** | **10** | **4.55** | **.005** | **3771** |
| R Putamen | 28 | -1 | 10 | 3.78 |  |  |
| **L Thalamus/Putamen** | **-19** | **-7** | **7** | **4.43** | **.008** | **3329** |
| Left Putamen | -23 | 11 | 10 | 3.90 |  |  |
| Left Putamen | -26 | -9 | 13 | 3.77 |  |  |
| **L Superior Temporal Pole** | **-40** | **21** | **-31** | **4.18** | **.017** | **2813** |
| L Superior Temporal Pole | -48 | 22 | -25 | 3.86 |  |  |
| L Middle Temporal Pole | -38 | 22 | -42 | 3.80 |  |  |
| *ii. DE>CE – No suprathreshold clusters* |  |  |  |  |  |  |
| *iii. CE>HE – No suprathreshold clusters* |  |  |  |  |  |  |
| *iv. HE>CE – No suprathreshold clusters* |  |  |  |  |  |  |
| *v. HE>DE* |  |  |  |  |  |  |
| **L Inferior Temporal Pole** | **-54** | **-20** | **-35** | **4.15** | **.022** | **2650** |
| L Inferior/Middle Temporal Pole | -62 | -22 | -33 | 4.03 |  |  |
| L Inferior Temporal Pole | -47 | -7 | -51 | 3.98 |  |  |
| *vi. DE>HE – No suprathreshold clusters* |  |  |  |  |  |  |

Bold face indicates peak voxels within cluster detected at $p < .001$ (uncorrected) at voxel level and FWE correction ( $p < .05$ ) at cluster level, with $k \geq 1000$ voxels.

### Post-hoc T-contrasts: RBM Results

CE>DE and HE>DE contrasts revealed significantly higher volumes for Right Putamen and Left Putamen ROIs for the CE (survived p < .05 FDR correction) and HE (uncorrected p < .05 only) groups (see Table 5).

**Table 5:** RBM T-contrast results.

| ROI | *p* value |  |
| --- | --- | --- |
|  | *uncorrected* | *FDR corrected* |
| ***i. CE > DE*** |  |  |
| R Putamen | < .001 | .003 |
| R Hippocampus | .023 | - |
| R Superior Occipital Gyrus | .024 | - |
| L Putamen | < .001 | .012 |
| L Post Central Gyrus | .002 | .028 |
| L Para Hippocampal Gyrus | .011 | - |
| L Supramarginal Gyrus | .019 | - |
| ***ii. DE > CE*** |  |  |
| R Lateral Orbital Gyrus | .035 | - |
| R Superior Orbital Gyrus | .035 | - |
| L Lateral Orbital Gyrus | .004 | - |
| ***iii. CE > HE*** |  |  |
| R Superior Occipital Gyrus | .022 | - |
| R Hippocampus | .024 | - |
| R Mid Occipital Gyrus | .038 | - |
| L Post Central Gyrus | .002 | .039 |
| L Superior Occipital Gyrus | .003 | .039 |
| L Mid Occipital Gyrus | .006 | - |
| L Hippocampus | .011 | - |
| L Lingual Gyrus | .027 | - |
| ***iv. HE > CE – Nil*** |  |  |
| ***v. HE > DE*** |  |  |
| R Putamen | .005 | - |
| L Putamen | .01 | - |
| L Inferior Temporal Gyrus | .02 | - |
| ***vi. DE>HE*** |  |  |
| R Lateral Orbital Gyrus | .048 | - |
| R Mid Orbital Gyrus | .019 | - |
| L Lateral Orbital Gyrus | .033 | - |

### Brain-Behaviour Bivariate Correlations

The pre-normalized GM volumes were extracted for the Left and Right Putamen which was identified in both VBM and RBM analysis. Correlations of either region with L2AoA or with L1 and L2 Self-rated Proficiency were not significant (*p* > .05).

## Discussion

The aim of this study was to explore whether relative language distance (RLD) induces any differences in structural capacity within the BLC network. To this end, exploratory VBM and RBM analysis were performed on three groups of bilinguals with comparable demographic and bilingual profiles but with varying and distinct levels of RLD of their spoken language pairs. After regressing out possible contributions of variables like age, education level, socio-economic status, TIV and L2AoA the results revealed significant differences amongst the groups in the bilateral putamen. An examination of the VBM contrast estimates at the right putamen revealed the effect to be largest for the hd-CE group and least for the ld-DE group, with id-HE lying in between. Although the classical whole-brain analysis detected a difference only in the right putaminal volumes, the more sensitive RBM method uncovered a bilateral putaminal difference as well. Post-hoc T-contrasts between the groups confirmed that both CE and HE groups had significantly greater grey matter volumes (GMV) in the bilateral putamen than the DE group (Tables 4 & 5). Moreover, these differences were independent of the differences in the L2AoA of the groups as well, and persisted regardless of L2AoA’s inclusion as a potential confound (Supplementary Tables S1 & S2). Based on these results it is suggested that the putaminal differences across groups, detected by both types of morphometric methods, may be due to the factor of varying RLD of their spoken language-pairs.

The observed differences in the Putamen reported in this study are not that surprising. The putamen is a critical part of the BLC network and has long been implicated in articulatory aspects (encoding and production) of phonological processing (e.g., Bohland & Guenther, 2006; Oberhuber et al., 2013; Rauschecker et al., 2008; Rosen et al., 2000; Riecker et al. 2002; Tettamanti et al., 2005; see also meta-analysis by Viñas-Guasch & Wu, 2017). Clinical evidence of impaired motor speech patterns after putaminal lesions (e.g., Speedie et al., 1993; Pickett et al., 1998; Robles et al., 2005) further emphasizes and attests to its critical role in affording coherent speech. Through its connections to the motor, pre-motor and prefrontal cortices (Lehericy et al, 2004a&b) the putamen aids articulatory control, which involves correct assemblage of phonemes and precise temporal sequencing of the required articulatory motor movements during speech production (Booth et al, 2007; Ullman, 2001). Considering that bilinguals would need this ability for not one, but two languages, articulatory control is arguably a vital component of language control for them. Several structural imaging studies examining bilingual brains confirm this by reporting structural changes in the putamen, specifically increased GM volumes, attributable to articulatory control demands (left Putamen: Abutalebi et al., 2013b and Berken et al., 2016; bilateral putamen: Burgaleta et al., 2015 and Pliatsikas et al., 2017). Because bilinguals, compared to their monolingual counterparts, must manage a significantly expanded articulatory/phonemic repertoire that spans both their languages and rely on the putamen for the same, the structure undergoes neuroplastic capacity increases. The finding of the present study is also in keeping with this line of inference. Among the three bilingual groups, the high-distance Cantonese-English speakers (CE) can be considered to have the most expanded and least overlapping phonemic repertoire due to the distinctness between their language pairs with respect to phonology. The Hindi-English (HE) bilinguals can be said to have more overlap between their languages than the CE group, but certainly not as much as the low-distance Dutch-English speakers (DE), whose languages have comparatively similar overlapping phonologies. Therefore, DE speakers have the smallest and most overlapping L1-L2 phonemic repertoires among the three groups. The articulatory control needs for managing L1-L2 phonemic repertoires with different extents of overlap/similarity would surely vary. Distant L1-L2 pairs with dissimilar L1 and L2 phonemic repertoires (like Cantonese and English) would likely require relatively more articulatory control owing to a greater variety of phonemes and distinct articulatory motor sequences than those language pairs whose respective phonemic repertoires are similar and overlapping (e.g., Dutch and English, comparatively similar phonemes and less unique articulatory sequences). Thus, the specific pattern of relative putaminal volumes (CE>HE>DE) corresponds to the progressively decreasing articulatory control needs generated by progressively decreasing L1-L2 RLD levels. Given that all three bilingual groups were fluent and experienced speakers of their language pairs who acquired both languages early on, they would have surely mastered the different articulatory/speech motor sequences necessary for speaking in their L1 and L2. Thus, they have all had sufficient experience in controlling speech and articulation in their respective spoken language pairs. Therefore, it is very much possible that the differing articulatory control needs induced by RLD differences, the only major experiential dimension that the bilingual groups varied on, has driven structural capacity changes in the putamen. Although articulatory control of speech involves several other neural regions as well (such as the frontal, motor cortices and the cerebellum), the evidence from the current study, along with those from previous studies in bilinguals involving the putamen, suggests that this region of the BLC network is a particularly susceptible locus for adaptive structural change. Overall, this finding also reinforces the notion that changes in the bilingual brain are merely a type of experience-dependant adaptation (Bialystok, 2017). As already outlined in the Adaptive Control Hypothesis (Green & Abutalebi, 2013), variation in bilingualism due to variation in the different experiential dimensions of bilingualism (such as proficiency, exposure, RLD etc.), gives rise to differing language control needs which the brain then accommodates via neuroplastic changes.

The results of the present exploratory study provide a positive indication of how the dimension of Relative Language Distance could influence bilingual brain structure, specifically that of the BLC network, after other major dimensions of bilingual experience are controlled for. Future studies employing other structural capacity measures such as shape analysis, cortical thickness and white matter connectivity analysis should be able to provide a clearer picture of the extent of RLD-mediated structural brain changes. This will further our understanding of how the bilingual brain is shaped by the linguistic experiences it is exposed to.

## Acknowledgements

The support and assistance of the technical staff at the University of Hong Kong’s 3T MRI Unit in MRI data collection is gratefully acknowledged.

## Footnotes

1 for evidence of bilingualism-associated *white* matter structural changes, see reviews by Bialystok, 2017; Li et al., 2014; Gold, 2015; Perani & Abutalebi, 2015

