## Supplementary Materials for "Exploring the effect of Relative Language Distance on Bilingual Brain Structure – a cross-sectional VBM study"

**Supplementary Table S1**: Whole brain VBM results for analysis without L2AoA, i.e. with only Education, SES, Age and TIV as potential confounds

| Region | *MNI co-ordinates (mm)* | | | *z*-score | Cluster *p* (FWE-corr) | *k* |
|---|---|---|---|---|---|---|
| | *x* | *y* | *z* | | | |
| **L Superior Temporal Pole** | **-41** | **23** | **-32** | **4.60** | **.007** | **2560** |
| L Mid Temporal Pole | -37 | 10 | -37 | 3.36 | | |
| L Mid Temporal Pole | -30 | 14 | -49 | 3.23 | | |
| **R cerebellum (Crus I)** | **55** | **-59** | **-28** | **4.57** | **.009** | **2382** |
| R cerebellum (Crus I) | 50 | -68 | -26 | 4.11 | | |
| R cerebellum (Crus I) | 55 | -64 | -40 | 3.41 | | |
| **R Putamen** | **23** | **2** | **7** | **4.36** | **<.001** | **4874** |
| R Putamen | 33 | -13 | -9 | 4.00 | | |
| R Putamen | 30 | -2 | 12 | 3.97 | | |

Table shows clusters with minimum of 1000 voxels per cluster > 8.0 mm apart. Bold Rows indicate peak voxels within cluster. Labels derived from AAL (Automatic Anatomical Labelling, Eickhoff et al., 2009)

**Supplementary Table S2**: RBM results for analysis without L2AoA, i.e. with only Education, SES, Age and TIV as potential confounds

| | *p* Value | |
|---|---|---|
| | uncorrected | FDR corrected |
| R Putamen | < .001 | 0.002 |
| L Putamen | < .001 | 0.01 |
| R Hippocampus | < .001 | - |
| L Post Central Gyrus | < .001 | - |
| L Superior Occipital Gyrus | 0.02 | - |
| L Hippocampus | 0.02 | - |
| L Middle Occipital Gyrus | 0.03 | - |
| L Para Hippocampal Gyrus | 0.03 | - |
| L Lateral Orbital Gyrus | 0.03 | - |

Table provides uncorrected and FDR corrected p-values for ROIs that significantly varied across groups. '-' indicates the ROI did not survive FDR correction.
